# The CIpP activator, TR-57, is highly effective as a single agent and in combination with venetoclax against CLL cells *in vitro*

**DOI:** 10.1101/2022.03.07.483345

**Authors:** Narjis Fatima, Yandong Shen, Kyle Crassini, Edwin J. Iwanowicz, Henk Lang, Donald S. Karanewsky, Richard I Christopherson, Stephen P Mulligan, O. Giles Best

## Abstract

Despite advances in treatment, a significant proportion of patients with chronic lymphocytic leukaemia (CLL) will relapse with drug-resistant disease.

Recent studies demonstrate that the imipridones ONC-201 and ONC-212 and the more potent TR-compounds are effective against a range of different cancers, including acute myeloid leukaemia and tumours of the brain, breast, and prostate. These drugs induce cell death through inhibition of mitochondrial function and activation of the mitochondrial protease, caseinolytic protease (CIpP), and the unfolded protein response (UPR).

Here we demonstrate that a drug in this class, TR-57, has efficacy as a single agent and is synergistic with venetoclax against CLL cells cultured under *in vitro* conditions that mimic the tumour microenvironment. The inhibitory effects of TR-57 on cell survival, proliferation and migration were irrespective of poor-risk features, including aberrations of TP53. Changes in protein expression suggest the mechanisms of action of TR-57 and its synergy with venetoclax involve activation of the UPR, inhibition of the AKT and ERK1/2 pathways and a pro-apoptotic shift in expression of proteins of the BCL-2 family.

The study suggests TR-57, as a single agent and in combination with venetoclax, may represent an effective treatment option for CLL, including for patients with poor-risk disease.

## Introduction

Chronic Lymphocytic leukaemia (CLL) is the most common form of adult leukaemia, accounting for approximately one third of all leukaemia’s diagnosed world-wide (2020). CLL is characterised by the clonal expansion of CD5-positive B-cells in the peripheral blood, bone marrow, lymph nodes and spleen (Gatt and Izraeli 2019). CLL cells in the peripheral circulation are predominantly quiescent, while cells within the lymph nodes and bone marrow proliferate in pseudofollicular structures known as proliferation centres (Herishanu, *et al* 2011). The regions within the lymph nodes and bone marrow populated by CLL cells are collectively known as the tumour microenvironment (TME); proliferation and survival of CLL cells in the TME is driven by the interaction of the leukemic cells with the ‘accessory’ cells and other immune cells that reside in these tissues and by a range of cytokines and growth factors produced by these cells (Caligaris-Cappio, *et al* 2014).

Understanding the mechanisms that drive the survival and proliferation of CLL cells in the tumour microenvironment has led to the development of effective treatment options for the disease (Fatima, *et al* 2020). Ibrutinib (Herman, *et al* 2011) and venetoclax (Roberts, *et al* 2016, Souers, *et al* 2013), which target the BTK and BCL-2 proteins, respectively, have proven very effective, in front-line and relapsed or refractory disease. However, relapse and drug resistance are common among patients treated with these newer agents, highlighting the need for the development of new treatment strategies.

To survive and proliferate under the nutrient-deprived and hypoxic conditions within the TME, cancer cells rely on pathways that are collectively known as the unfolded protein response (UPR) (Wang, *et al* 2009). Recent studies suggest that targeting components of the UPR may represent an effective treatment approach, resulting in apoptosis in a variety of cancers (McConkey 2017). In healthy cells, the UPR functions to limit protein translation and cell proliferation in response to a variety of cellular stresses and may ultimately trigger apoptosis. However, the proliferation and survival of cancer cells is dependent on their ability to adapt to the stress associated with rapid cellular growth and limited nutrient and oxygen availability. Several key proteins and pathways have been identified that regulate the UPR. Endoplasmic reticulum (ER) stress activates the Grp78 protein, which in turn activates protein kinase RNA-like endoplasmic reticulum kinase (PERK), inositol-requiring enzyme 1 (IRE1) and activating transcription factor 6 (ATF6) (Lee 2005). PERK phosphorylates eIF2α, which generally attenuates protein synthesis. However, the UPR also increases translation of certain proteins through activity of transcription factors, including ATF4 (Wang, *et al* 2009).

Several recent studies have demonstrated that a class of drugs known as imipridones have efficacy against a range of cancers (Allen, *et al* 2015, Prabhu, *et al* 2018), including acute myeloid leukaemia (AML) (Cole, *et al* 2015). The mechanisms of action of the imipridones include inhibition of mitochondrial function (Greer, *et al* 2018) and activation of the mitochondrial protease, caseinolytic protease proteolytic subunit (CIpP) (Graves, *et al* 2019, Ishizawa, *et al* 2019, Wang and Dougan 2019). Paradoxically, ClpP overexpression has been observed in several cancers that are sensitive to the imipridones, including AML (Cole, *et al* 2015), prostate and breast (Seo, *et al* 2016).

In our recent study, we demonstrated that the imipridone ONC-212 (TR-31) induces expression of ClpP, inhibits key signalling pathways downstream of the BCR and induces a pro-apoptotic shift in the expression of proteins of the BCL-2 family in CLL cells, under *in vitro* conditions that mimic the TME (Fatima, *et al* 2021). These data provided the rationale for the current study, in which we investigated the effects of TR-57 towards CLL cells under conditions that mimic the pro-survival effects of the TME. Based on our previous study and on the role of MCL-1 in determining venetoclax sensitivity (Gong, *et al* 2016), we also examined the possibility that synergy between TR-57 and the BH3-mimetic may make this an effective treatment option for patients with CLL.

## Materials and methods

### Patient samples

Samples were collected from CLL patients managed at Royal North Shore Hospital, Sydney following informed, written consent. CLL was diagnosed according to the international workshop on CLL (iwCLL) guidelines (Hallek, *et al* 2008). Peripheral blood mononuclear cells (PBMCs) were isolated from whole blood by ficoll-density gradient centrifugation. All patient samples contained >85% CD5^+^/CD19^+^ (CLL) cells, as determined by flow cytometry (data not shown). Cells were cryopreserved in foetal calf serum (FCS), containing 10% dimethyl sulphoxide (DMSO). ATM/TP53 function and ZAP-70 and CD38 expression were assessed by flow cytometry, using previously published methods (Best, *et al* 2008, Orchard, *et al* 2004). Category 1 and 3 ATM/TP53 dysfunction are consistent with mutations in *TP53* (Best, *et al* 2008) and the presence of a small TP53 dysfunctional clone (Tracy, *et al* 2017), respectively. Deletions of the 17p13 and 11q23 chromosomal regions, which encompass *TP53* and *ATM*, respectively, were detected by fluorescence *in-situ* hybridisation (FISH).

### Cell culture

Primary cells and cell lines were cultured in RPMI-1640 (Thermo Fisher Scientific, Waltham, MA, USA) medium supplemented with 10% FCS, 2 mM L-glutamine and 1% penicillin/streptomycin (complete medium). The OSU-CLL cell line was obtained under a materials transfer agreement with at Ohio State University. Details of the derivation and characterisation of the OSU-CLL line are described in (Hertlein, *et al* 2013).

Primary CLL cells were seeded at 1 × 10^5^ or 2 × 10^6^ cells per well of a 96- or 24-well plate, respectively. Mouse L-fibroblasts, expressing the human CD40-ligand (CD40L-fibroblasts) were seeded at a density of 250 cells/µL one day prior to co-culture with primary CLL cells.

### Generation of a TP53 knock-out OSU-CLL cell line

Knock-out of the *TP53* gene in the OSU-CLL cell line was achieved using a lentiviral CRISPR-Cas9 technology developed at the Walter and Eliza Hall Institute (WEHI), Melbourne, Australia, with methods described by Aubrey *et al*. (Aubrey, *et al* 2015). Sanger sequencing and immunoblotting were performed to confirm complete knockout of the *TP53* gene and protein. Flow cytometry was used to confirm that the processes involved in the generation of the OSU-CLL*TP53*ko line had no effect on CD5/CD19 expression (data not shown).

### Assessment of cell viability

Primary CLL cells, the OSU-CLL cell lines and B-cells isolated from healthy donors were incubated with TR-57 and venetoclax at the doses and times indicated. Cell viability was assessed by flow cytometry on an LSR Fortessa instrument (Becton Dickinson, Franklin Lakes, NJ, USA) by staining cells for 20 min at 37°C with 0.05 µM of the mitochondrial membrane potential dye, 1,1’,3,3,3’,3’-hexamethylindodicarbocyanine iodide (DiIC_1_(5); ThermoFisher Scientific) and 10 µM propidium iodide (PI; Sigma-Aldrich, St Louis, MI, USA). Drug concentrations that induced a 50% decrease in cell viability compared to untreated controls (IC50 values) were calculated using GraphPad Prism software (GraphPad Software, San Diego, CA, USA).

The cytotoxic effects of TR-57 and venetoclax, alone and in combination, towards CLL cells and autologous T-cells were assessed by flow cytometry using antibodies against CD5 and CD19 and the viability dyes, DiIC_1_(5) and PI. The proportions of viable CLL (CD5^+^/CD19^+^) and T-cells (CD5^+^/CD19^-^) were determined before and after drug treatment.

### Assessment of drug synergy

TR-57 and venetoclax were combined at ratios based on their IC50 values as single agents (Supplementary Table 1). CompuSyn software (www.combosyn.com) was used to calculate combination indices (CI) for the drugs against the cell lines and primary CLL cells. CI values of <1, 1 and >1 are considered indicative of synergy, additivity and antagonism, respectively.

### Cell cycle and proliferation

Proliferation of the OSU-CLL and OSU-CLL-*TP53*ko cells was assessed using the amine dye, carboxyfluorescein succinimidyl ester (CFSE; Sigma-Aldrich). Cells were stained with 2 µM CFSE and cultured overnight prior to treatment with the IC25 doses of TR-57 and venetoclax, alone or in combination. The rate of decay of the mean fluorescence intensity (MFI) of CFSE at 24, 48, and 72 h was analysed by flow cytometry and used to calculate the rate of proliferation.

The effects of the drugs on the cell cycle distribution of primary CLL cells was assessed by first stimulating peripheral blood-derived CLL cells using 2 µM of the CpG-oligonucleotide, Dsp30 (Integrated DNA Technologies, Coralville, IO, USA), and 200 U/ml interleukin 2 (IL-2; Peprotech, Rocky Hill, NJ, USA) for 4 days. Stimulated CLL cells were treated with TR-57 and venetoclax, alone or in combination, for a further 48 h in the presence of Dsp30/IL2, then harvested into ice cold 50% ethanol for at least 24 h at -20°C. Samples were stained with 40 µg/ml PI, 100 µg/ml RNase and 0.1% Triton in PBS at 37°C for 30 min. Data was acquired by flow cytometry and analysed using ModFit software (Verity Software House, Topsham, ME, USA). Data are expressed as the proportion of CLL cells in the S and G2/M phases.

### Integrin and cytokine-receptor expression and cell migration

Primary CLL cells in co-culture with CD40L-fibroblasts were treated with 100 nM TR-57 or 10 nM venetoclax or the drugs in combination for 24 h. The cells were then harvested and stained with phycoerythrin (PE) conjugated antibodies to CD49d or CXCR4 (Biolegend, San Diego, CA, USA). The expression of both proteins on viable cells was assessed by flow cytometry. Results are expressed as the fold-change in the mean fluorescence intensity (MFI) of expression relative to expression on untreated cells.

The effects of the drugs on the migratory capacity of primary CLL cells was assessed using the CXCR4 ligand, SDF-1α, as a chemoattractant. Primary CLL cells were treated for 24 h with TR-57 and venetoclax, alone or in combination. Cell viability was assessed by trypan blue exclusion and 3 × 10^6^ viable cells from each condition were loaded into the upper chambers of Transwell culture inserts with a 0.5 µm pore size (Merck, Burlington, MA, USA). Cells were allowed to migrate towards the lower well containing medium, with or without 200 ng/ml SDF1α, for 3 h. Viable cells harvested from the lower well were identified by staining with DiIC_1_(5) and PI and enumerated by flow cytometry with an acquisition time of 120s. Data are expressed as a fold-change in the number of viable CLL cells that migrated through the insert, relative to untreated controls.

### Immunoblotting

Primary CLL cells cultured in medium alone or in co-culture with CD40L-fibroblasts were treated with TR-57 (500 nM) and venetoclax (50 nM), alone or in combination for 24 h. Viability of the CLL cells was assessed by trypan blue exclusion prior to cell lysis to exclude the possibility that the observed effects were due to differences in the proportion of dead cells present. Cells were then washed with PBS and lysed in radioimmunoprecipitation assay buffer (RIPA; 50 mM sodium chloride, 1.0% Triton X-100, 0.5% sodium deoxycholate, 0.1% sodium dodecyl sulphate, 50 mM Tris-HCl, pH 8.0) containing a cocktail of phosphatase and protease inhibitors (MSSafe; Sigma Aldrich, St Louis, MI). Methods used for immunoblotting and data analysis were as described in our previous study (Shen, *et al* 2019).

### Statistics

All statistical analyses were conducted using the students T-test function of GraphPad Prism software. P-values of < 0.05 were considered significant.

## Results

### TR-57 is cytotoxic towards CLL cells cultured under in vitro conditions that mimic the tumour microenvironment

TR-57 was effective at inducing apoptosis of primary CLL cells (n = 15 patients; CLL# 1,2,4,5,7,12,13,15-19,20-22) cultured in medium alone or with CD40L-expressing fibroblasts (Figure 1A). However, CLL cells cultured in medium alone were significantly more sensitive to TR-57 than cells in co-culture with CD40L-fibroblasts. The IC50 values for TR-57 were 38.0 ± 1.38 nM in medium compared to 287 ± 50.5 nM against CLL cells in stromal co-culture (Supplementary Table 1). No significant difference (P = 0.70) in sensitivity to TR-57 was observed between samples with (n = 5; CLL# 2,7,15,19,21) or without (n = 8; CLL# 1,4,5,16,17,18,20,22) TP53 aberrations (Figure 1B). Similarly, no association between ZAP-70 or CD38 expression and response of CLL cells to TR-57 was apparent.

**Figure 1.**
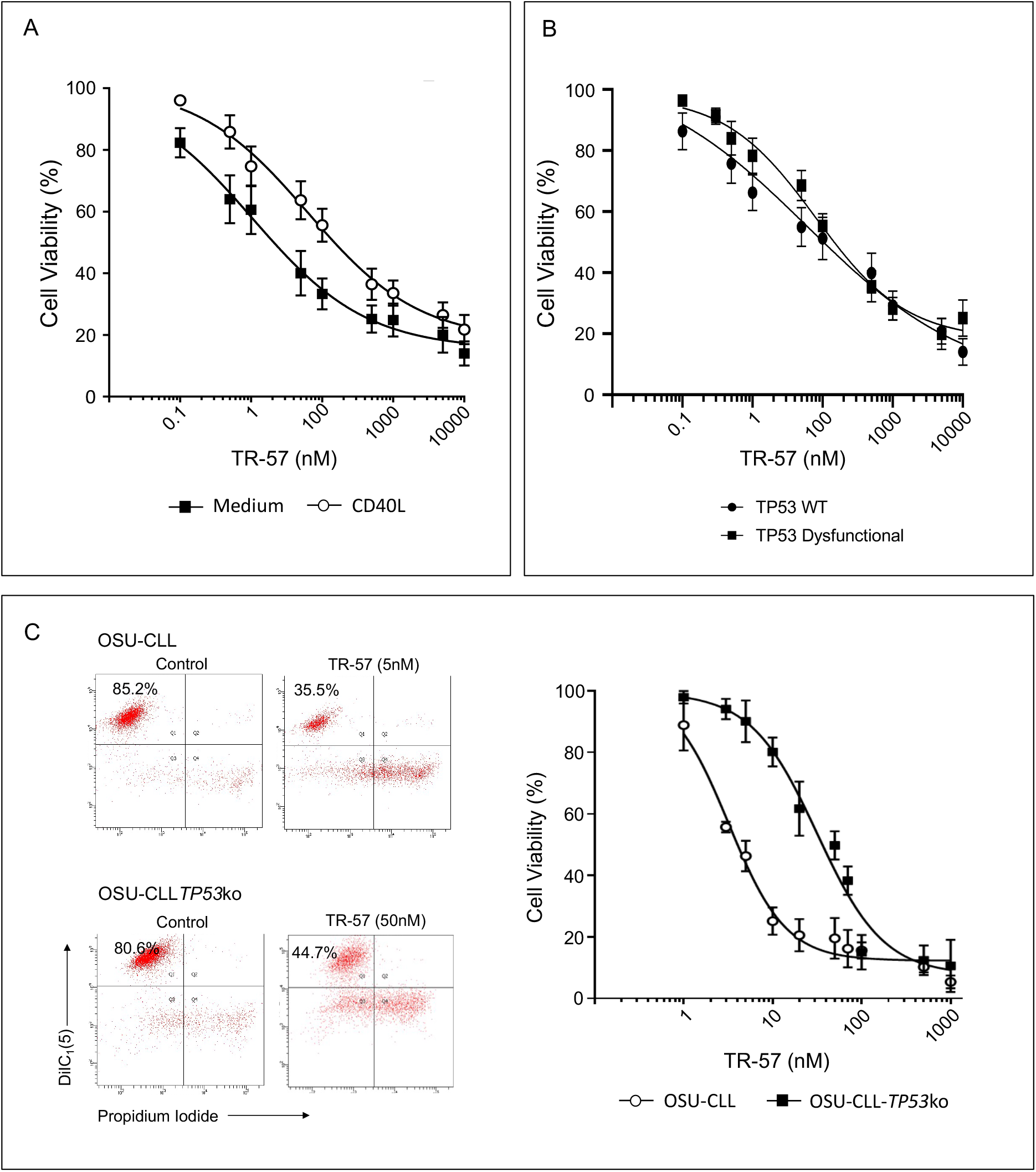
TR-57 induces apoptosis in primary CLL cells and in the OSU-CLL and OSU-CLL-*TP53*ko cell lines. CLL cell viability was assessed by flow cytometry using DiIC_1_(5) and propidium iodide. Viability is expressed relative to vehicle-treated controls and error bars represent the standard deviation. (A) Primary CLL cells (n = 15 patients), cultured in medium alone or in co-culture with CD40L-fibroblasts, were treated with TR-57 for 48 h. (B) Dose response analysis of TR-57 against TP53 wild-type (n = 10) and TP53 dysfunctional (n = 5) CLL patient samples following 48h treatment in co-culture with CD40L-fibroblasts. (C) OSU-CLL and OSU-CLL-*TP53*ko cell lines were treated with TR-57 for 48 h. Data are the mean of 4 biological replicates.

### TR-57 is cytotoxic towards TP53-deficient CLL cells

To further examine the effects of TR-57 on TP53-deficient CLL cells, we generated an OSU-CLL cell line in which *TP53* was knocked-out using the CRISPR-Cas9 technology (OSU-CLL-*TP53*ko). OSU-CLL cells transfected with CRISPR-Cas9, but not activated with doxycycline, were used as a control for possible effects of the transfection processes. No significant difference (P = 0.25) in the sensitivity of these control cells and OSU-CLL cells to TR-57 was observed (Supplementary Figure 1).

TR-57 was cytotoxic towards OSU-CLL and OSU-CLL-*TP53*ko cells in a dose-dependent manner, however the IC50 for TR-57 was significantly (P = 0.004) higher against the *TP53*ko than wild-type line (Figure 1B). The IC50 values were 3.98 ± 1.03 nM and 90.71 ± 3.51 nM, for the OSU-CLL and OSU-CLL-*TP53*ko lines, respectively (Supplementary Table 1).

### TR-57 and venetoclax are synergistic against primary CLL cells and OSU-CLL and OSU-CLL-TP53ko cells

Next, we explored the effects of combining TR-57 with venetoclax against primary CLL cells co-cultured with CD40L-expressing fibroblasts, including 5 samples from patients with ATM/TP53 deletions or dysfunction (CLL# 2,7,15,19,21; Table 1), and the OSU-CLL and OSU-CLL-*TP53*ko lines. TR-57 and venetoclax in combination had a significantly greater effect on cell viability than either drug alone against primary CLL cells (Figure 2A) and the OSU-CLL (Figure 2B) and OSU-CLL-*TP53*ko (Figure 2C) lines. Combination indices (CI) at a fractional effect of 0.5 (50% cell killing) for primary CLL cells, OSU-CLL and OSU-CLL-*TP53*ko cells, were 0.13, 0.10 and 0.10 respectively (Supplementary Table 1). CI values across the range of fractional effects assessed are shown in Supplementary Table 2.

**Table 1.**
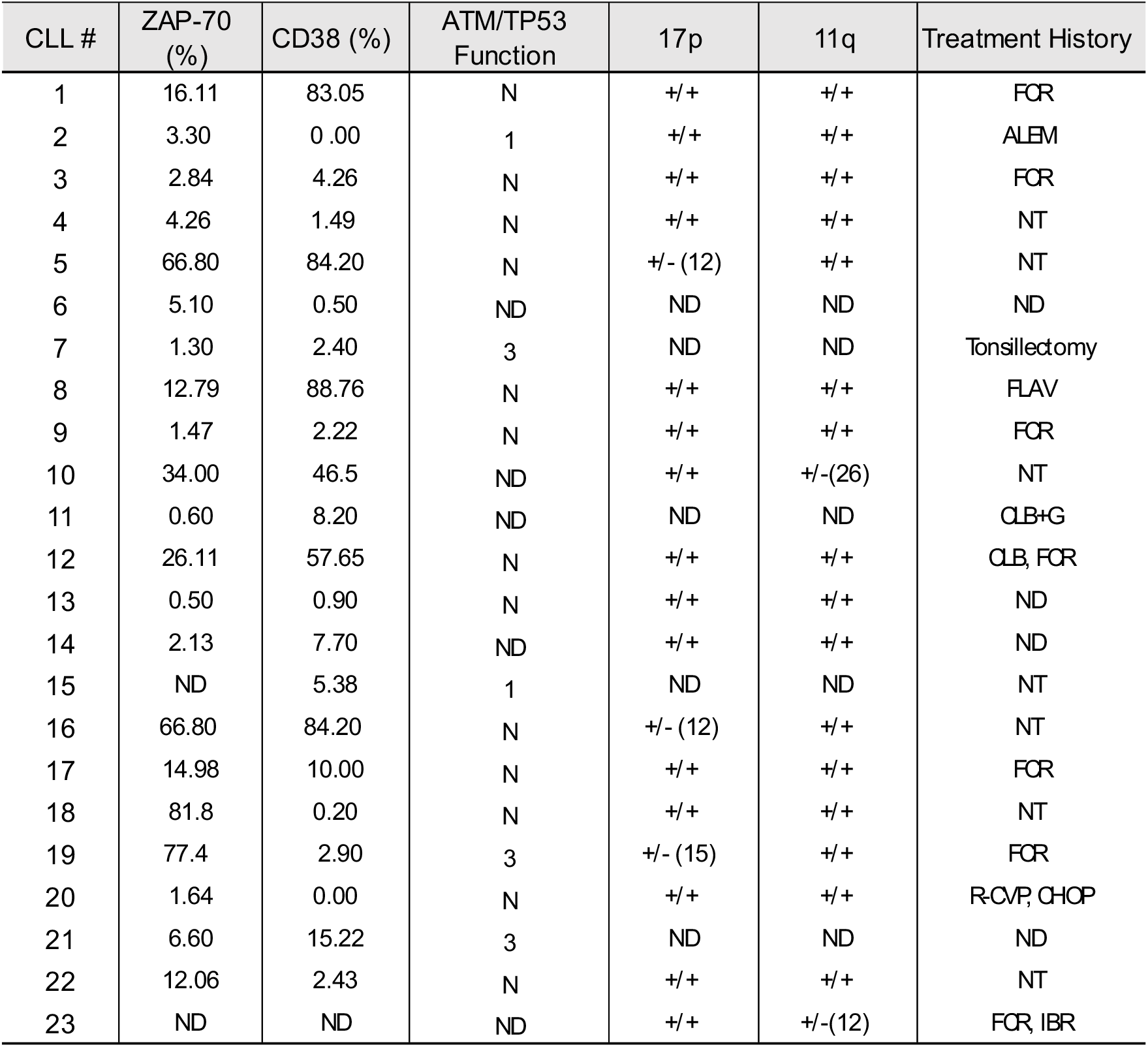
Details of CLL patient samples studied. CLL patient samples were assessed for ZAP-70 and CD38 expression and TP53 functional status by flow cytometry. Cut-offs for ZAP-70 and CD38 positivity were 10 and 30%, respectively. Patients were categorised based on TP53 functional status as either N – normal, 1 – TP53 dysfunction or 3 – small TP53 dysfunctional clone. Deletions of chromosomes 17p13 and 11q23 were assessed by fluorescence *in-situ* hybridisation (FISH); +/+ and +/- denote no deletion and a heterozygous deletion of one allele, respectively. Clone size of the deletions are shown in parentheses. Abbreviations for treatment history are as follows: FCR – fludarabine, cyclophosphamide, rituximab; ALEM – alemtuzumab; FLAV – flavopiridol; G – obinutuzumab; CLB – chlorambucil; R-CVP – rituximab, cyclophosphamide, vincristine and prednisone; CHOP – cyclophosphamide, doxorubicin, vincristine and prednisone. NT – no treatment. ND indicates no data.

**Figure 2.**
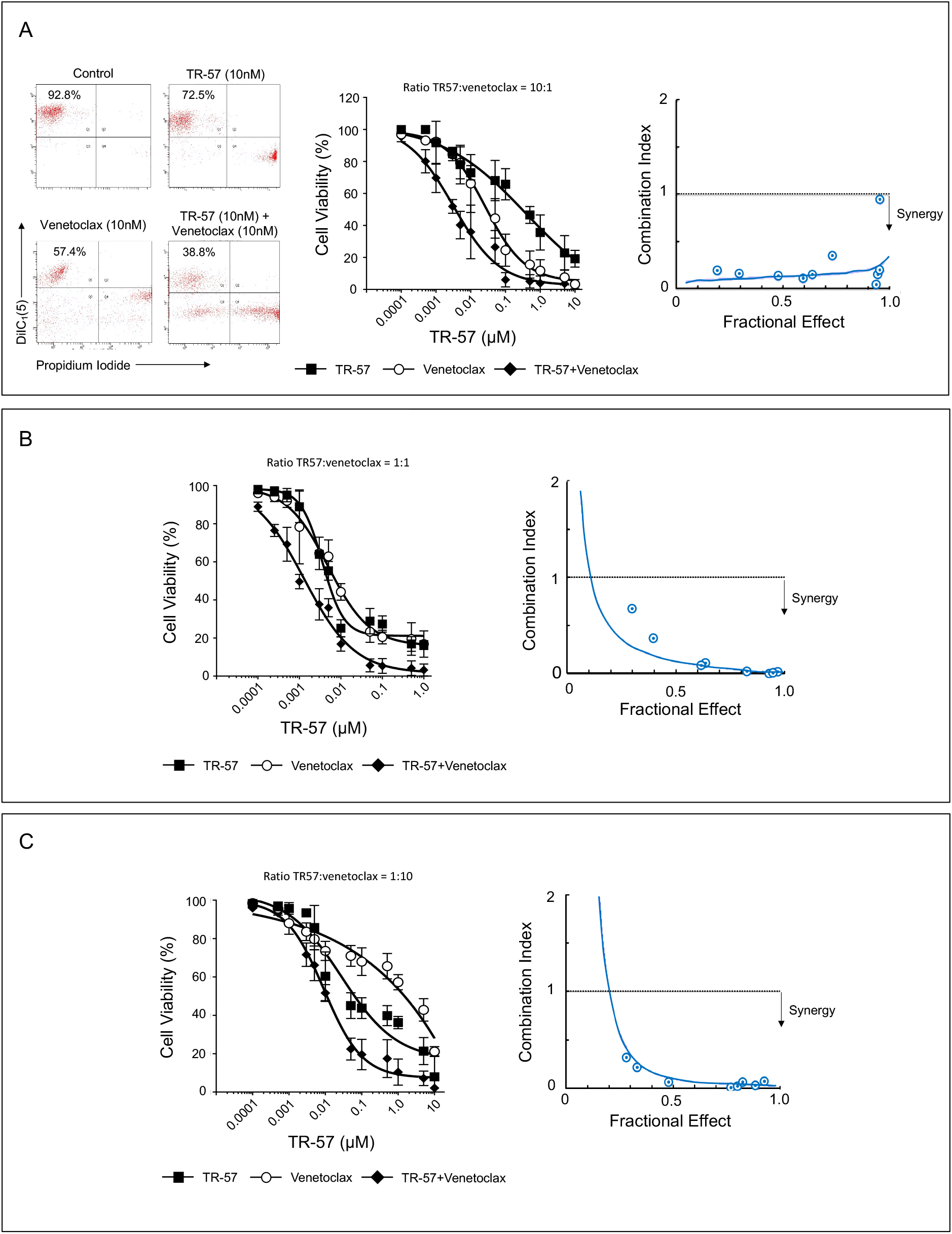
TR-57 and venetoclax are synergistic in their cytotoxic effects towards CLL cells. Cell viability was assessed by flow cytometry using DiIC_1_(5) and propidium iodide following treatment with TR-57 and venetoclax as single agents and in combination. Drugs were combined at the fixed ratios indicated, based on their IC50 values as single agents. Synergy was assessed by calculating combination indices (CI) at a range of fractional effects using the CompuSyn software. CI values of >1, 1 and <1 are indicative of antagonism, additivity and synergy, respectively. Dose-response analyses and assessment of synergy in (A) CLL patient samples (n = 15). Representative data from one CLL patient sample is shown. (B) OSU-CLL and (C) OSU-CLL-*TP53*ko cell lines. Error bars represent the standard deviation. Data in (B) and (C) are the mean of 3 biological replicates.

The cytotoxic effects of TR-57 and venetoclax, alone and in combination, were also examined against PBMC’s from healthy donors and against T-cells in PBMC fractions from CLL patients (Supplementary Figure 2B). Healthy PBMC’s were significantly less sensitive to both drugs as single agents (P = 0.0001) or in combination (P = 0.001). IC50 values for TR-57, venetoclax and the drugs in combination are shown in Supplementary Table 1. TR-57 and venetoclax, alone (P = 0.04 and P = 0.03, respectively) and in combination (P = 0.01) significantly reduced the proportion of viable CLL cells, with a concomitant increase in the proportion of viable T-cells, in PBMC fractions from CLL patients (n=5, CLL# 2,8,9,14,21; supplementary Figure 2C).

### TR-57 and venetoclax in combination arrest the proliferation and cell cycle progression of CLL cells

TR-57 and venetoclax, alone or in combination, significantly decreased the rate of proliferation of both the OSU-CLL and OSU-CLL-*TP53*ko cells (Figure 3A). After 72h, we observed a significant (P = 0.002 and 0.004 in the OSU-CLL and OSU-CLL*TP53*ko lines, respectively) reduction in the degree of cell proliferation following treatment with TR-57 and venetoclax in combination, compared to cells treated with either drug as a single agent. Stimulation of primary CLL cells from 7 patients (CLL# 3,6,7,10,11,19,23) with Dsp30/IL2 induced a significant (P = 0.005) increase in the percentage of cells in G2/M/S-phase compared to cells cultured in medium alone (Figure 3B). TR-57, alone (P = 0.03) and in combination with venetoclax (P = 0.009), significantly reduced the proportion of cells in the G2/M/S-phases.

**Figure 3.**
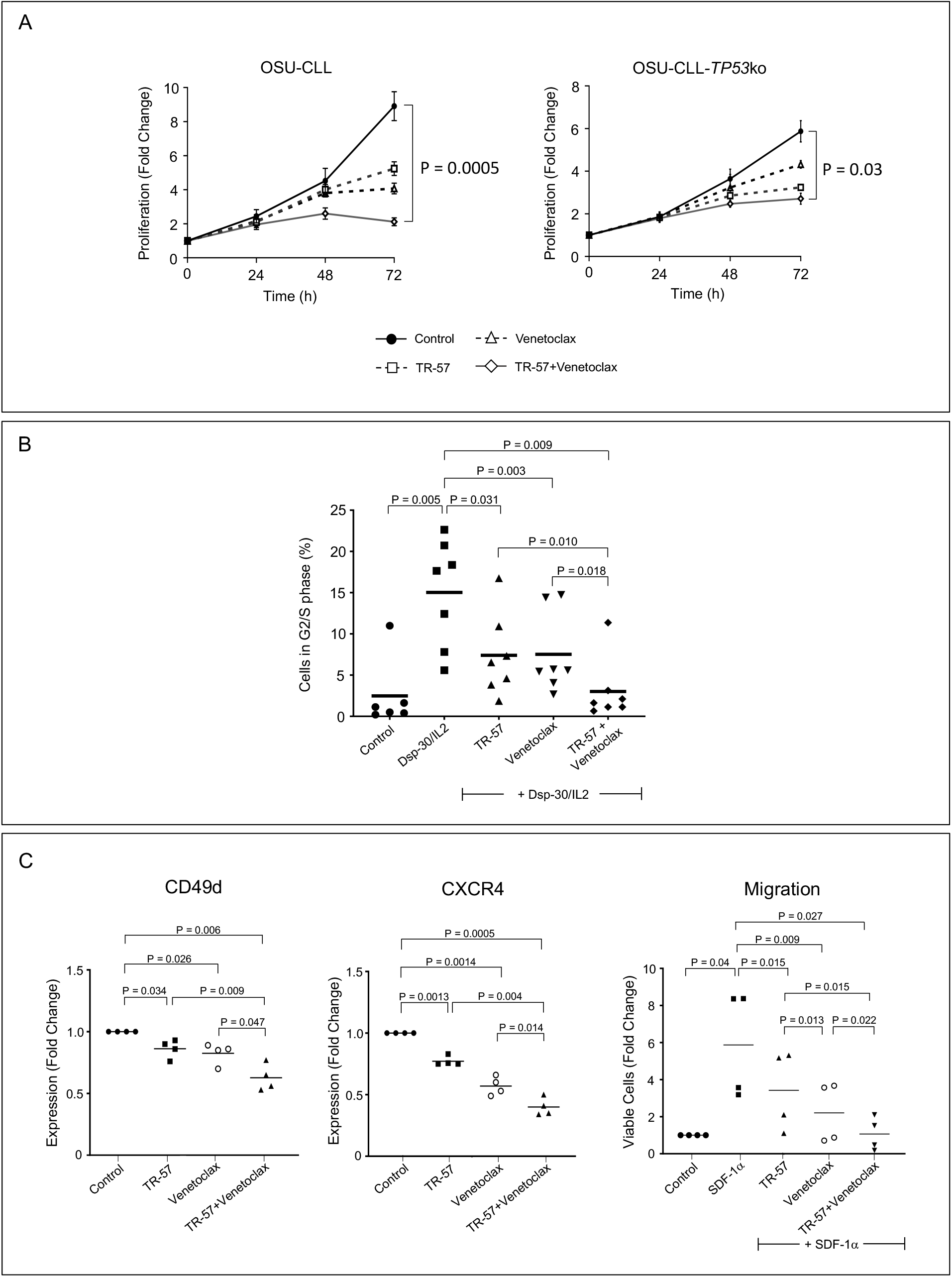
TR-57 and venetoclax reduce the proliferative and migratory capacity of CLL cells. (A) Proliferation of the OSU-CLL and OSU-CLL-*TP53*ko cells was assessed using flow cytometry in cells stained with CFSE. Cells were treated with TR-57 and venetoclax, alone or in combination for 0 – 72 h (2 and 2 nM and 25 and 250 nM for TR-57 and venetoclax against the OSU-CLL and OSU-CLL*TP53*ko lines, respectively). (B) Cell cycle analysis by flow cytometry of primary CLL cells (n = 7 patients) following culture in medium alone or stimulation with Dsp30/IL2. The proportion of CLL cells stimulated into S or G2/M phase and the effects of TR-57 and venetoclax, alone or in combination, on the stimulation was assessed. Representative histograms from one patient sample are shown. Data were analysed using ModFit software. (C) The effects of TR-57 and venetoclax, alone or in combination, on the expression of CD49d (left) and CXCR4 (centre) and on the migratory capacity (right) of primary CLL cells was assessed by flow cytometry. Data are expressed as fold-changes relative to cells cultured in medium alone (control).

### TR-57 and venetoclax in combination decrease expression of CD49d and CXCR4 and attenuate the migratory capacity of CLL cells

The chemokine receptor CXCR4 and the integrin CD49d play important roles in the homing and retention of CLL cells within the TME (Burger, *et al* 1999, Burger and Kipps 2002, Kriston, *et al* 2018). TR-57 and venetoclax treatment resulted in a significant decrease in the expression of CD49d (P = 0.03 and 0.02, respectively) and CXCR4 (P = 0.001 for both drugs), compared to expression on vehicle-treated cells (CLL# 1,14,15,19; Figure 3C). The expression of both proteins was significantly lower (P = 0.008 and 0.0004 for CD49d and CXCR4, respectively) on cells treated with both drugs than on cells treated with either drug alone.

The functional consequence of CXCR4 downregulation was examined by assessing the migration of primary CLL cells towards the CXCR4 ligand, SDF-1α. The number of viable CLL cells that migrated across the permeable support was significantly (P = 0.04) higher in the presence of SDF-1α than in medium alone (Figure 3C, right). Consistent with the effects of the drugs on expression of CXCR4 (Figure 3C, centre), the drugs significantly reduced the number of viable CLL cells that migrated under the influence of SDF-1α (P = 0.01 for both drugs). The migration of CLL cells treated with TR-57 and venetoclax in combination was significantly (P = 0.02) lower compared to cells treated with either drug as a single agent. Treatment with the drugs in combination resulted in a complete attenuation of SDF-1α–induced migration, relative to cells in medium alone (P = 0.14).

### TR-57 and venetoclax in combination induce expression of CIpP and ATF4 and decrease HSP60 in CLL cells

Next, changes in protein expression following treatment with TR-57 and venetoclax were examined by immunoblotting. All fold changes in expression are the mean of all samples assessed. Co-culture of CLL patient samples (n = 4; CLL# 1,7,14,19) with CD40L-expressing fibroblasts decreased expression of CIpP and increased expression of ATF4 and Grp78 in all samples, relative to expression in cells cultured in medium alone (Figure 4A). CD40L-fibroblast co-culture also increased expression of the molecular chaperone HSP60 in 3 of the 4 samples. Treatment of CLL cells in CD40L-fibroblast co-culture with TR-57 increased expression of CIpP (3 samples: 1.60-fold) and ATF4 (4 samples: 7.02-fold) and decreased expression of HSP60 (4 samples: 0.75-fold). No consistent effect of TR-57 on Grp78 expression was observed.

**Figure 4.**
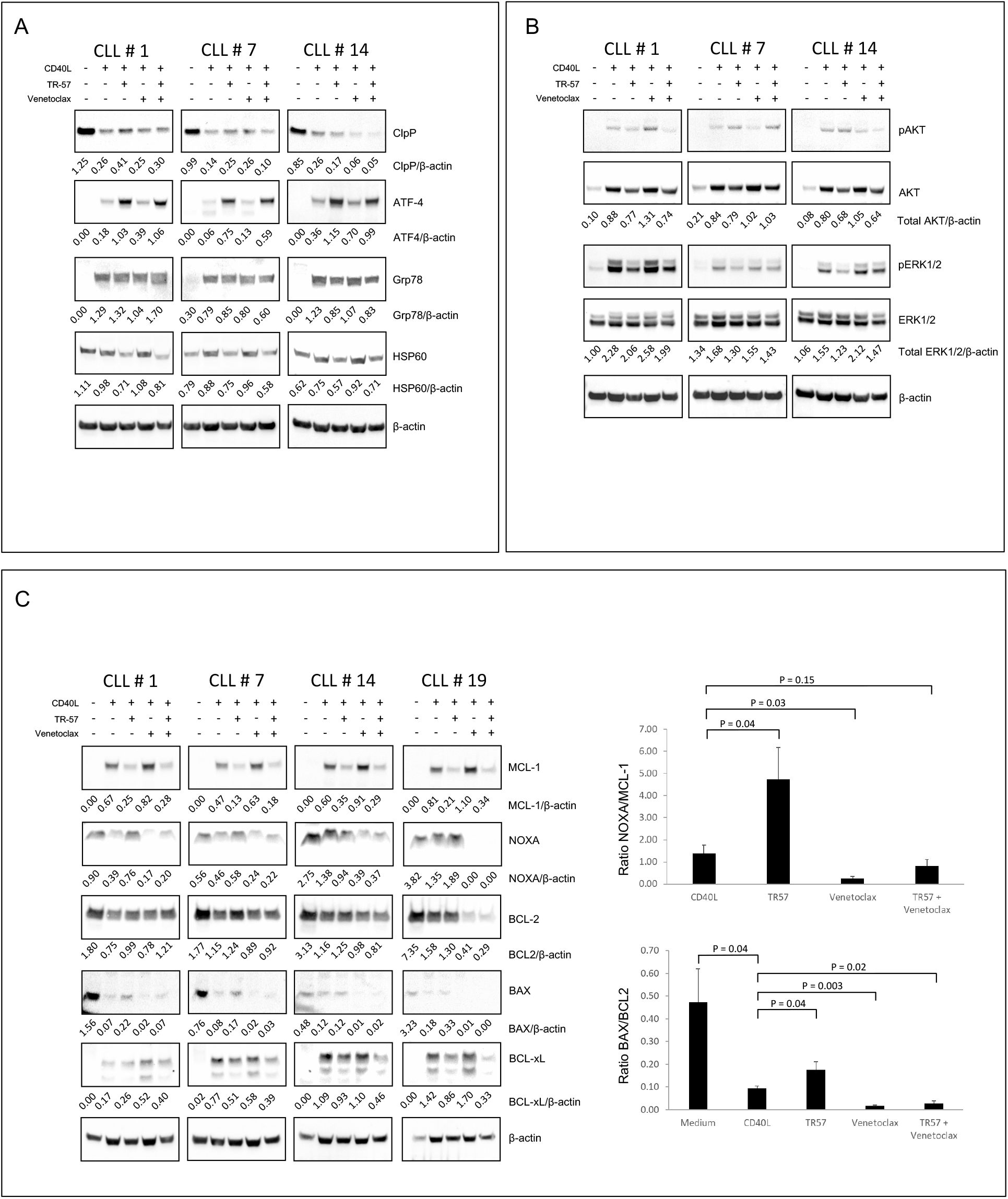
TR-57, alone and in combination with venetoclax, induces changes in proteins involved in the UPR, inhibits AKT and ERK1/2-MAPK signalling and induces a pro-apoptotic shift in BCL-2 family proteins. Immunoblotting of CLL patient samples examining changes in (A) UPR-related proteins, CIpP and HSP60, (B) AKT and ERK1/2-MAPK signalling and (C) BCL-2 family proteins. Primary CLL cells were cultured in medium alone or with CD40L-fibroblasts and were treated either with vehicle, 500 nM TR-57 or 50 nM venetoclax, as single agents or in combination, for 24 h. The ratios of proteins to β-actin shown under the immunoblots or pro- to anti-apoptotic BCL-2 proteins shown in the histograms (C) were calculated using ImageJ software.

Venetoclax had variable effects on the expression of CIpP, but increased expression of ATF4 in all 4 samples (1.74-fold), albeit to a lesser extent than TR-57. Decreased levels of Grp78 were also observed in 2/4 samples following treatment with venetoclax. In cells treated with TR-57 and venetoclax in combination, we observed decreased expression of CIpP in 3/4 samples (Figure 4A: 0.51-fold). In all 4 samples, ATF4 levels were increased relative to CLL cells in CD40L-fibroblast co-culture (5.17-fold). Expression of Grp78 and HSP60 was decreased in 3/4 (0.88-fold) and 4/4 (0.79-fold) samples, respectively following treatment with the drugs in combination, relative to expression in CLL cells in co-culture with CD40L-fibroblasts.

### TR-57 and venetoclax in combination reduce phosphorylation of AKT and ERK1/2-MAPK in CLL cells

TR-57 reduced expression of total (phosphorylated and pan-protein) AKT and ERK1/2-MAPK by 0.89 and 0.82-fold, respectively, in 4 CLL patient samples relative to CD40L-fibroblast co-cultured controls (Figure 4B). In contrast, increased expression of total AKT (1.34-fold) and ERK-1/2 (1.14-fold) protein was observed in cells treated with venetoclax. TR-57 and venetoclax in combination reduced total expression of AKT (3/4 samples) and ERK1/2-MAPK (4/4 samples).

### TR-57 and venetoclax in combination induce a pro-apoptotic shift in the expression of BCL2-family proteins

TR-57 treatment decreased stroma-induced expression of MCL-1 (0.37-fold) and BCL-xL (0.92-fold) in all 4 samples assessed but only decreased expression of BCL-2 in 1 sample; an increase in expression was observed in the remaining 3 samples (Figure 4C). In 3/4 samples, increased expression of the pro-apoptotic NOXA (1.32-fold) and BAX (2.02-fold) proteins was also observed. A significant increase in both the NOXA/MCL-1 and BAX/BCL2 ratios was observed in cells treated with TR-57, compared to controls (Figure 4C, histograms). Venetoclax decreased expression of BCL-2 in 3/4 samples, but interestingly increased expression of MCL-1 and BCL-xL in 4 and 2 samples, respectively. Venetoclax reduced levels of both NOXA and BAX, which resulted in a decrease in the NOXA/MCL-1 and BAX/BCL-2 ratios, relative to control cells. Treatment with TR-57 and venetoclax in combination resulted in decreased expression of MCL-1 (0.43-fold), BCL-2 (0.83-fold) and BCL-xL (0.89-fold) compared to untreated controls in 3 or more of the samples.

## Discussion

Despite high response rates and improved outcomes associated with BTK and BCL-2-targeted therapies, a significant proportion of patients with CLL experience disease relapse, due in part to the evolution of drug-resistant disease clones. CLL is still widely considered an incurable disease and there remains a need for the on-going development of novel treatment strategies with a curative intent.

Studies suggest drugs that target the unfolded protein response (UPR) and upregulate caseinolytic protease (ClpP) activity may be an effective treatment option for several cancers (Graves, *et al* 2019). Our previous study (Fatima, *et al* 2021) and development of the potent CIpP-activating, TR-compounds prompted the current study, in which we examined the effects of TR-57 against CLL cells, as a single agent and in combination with venetoclax.

TR-57 had both cytotoxic (Figure 1) and cytostatic (Figure 3) effects against primary CLL cells and CLL cell lines, including a line in which *TP53* was knocked-out using CrispR-Cas9 (OSU-CLL-*TP53*ko). In comparison to data from our previous study (Fatima, *et al* 2021), TR-57 was significantly (P = 0.0001) more cytotoxic towards CLL cells than ONC-212, with IC50 values of 287 ± 50.5 nM and 404 +/- 70.6 nM, respectively against primary CLL cells in co-culture with CD40L-fibroblasts. The cytotoxic effects of TR-57 against CLL cells under *in vitro* conditions that mimic the tumour microenvironment were consistent with the ability of the drug to counter the supportive effects of the stromal cells and induce a pro-apoptotic shift in expression of MCL-1, BCL-xL, NOXA and BAX and phosphorylation of AKT and ERK1/2-MAPK (Figure 4). Importantly, TR-57 was effective against primary CLL cells from patients with ATM or TP53 dysfunction (Figure 1A) and OSU-CLL-*TP53*ko cells (Figure 1B). The significant reduction in sensitivity of CLL cells to TR-57 when in co-culture with stromal cells may be associated with decreased expression of CIpP, NOXA and BAX and increased expression of AKT, ERK1/2-MAPK, MCL-1 and BCL-xL (Figure 4), which collectively may increase the ‘apoptotic threshold’ in these cells. The reduced sensitivity of the OSU-CLL*TP53*ko line to venetoclax may also be related to the role of TP53 in the induction of the pro-apoptotic BAX protein (Chipuk, *et al* 2004); the BAX/BCL-2 ratio in the OSU-CLL*TP53*ko cells was approximately 1/3 of that observed in the wild-type line following treatment (Supplementary Figure 3). While the OSU-CLL*TP53*ko line represents a model for complete absence of TP53, it is widely recognised that *TP53* mutations often result in a protein with some residual activity or a ‘gain of function’, which may explain why *TP53*ko and not TP53 dysfunction results in a significant decrease in sensitivity to TR-57. Despite the difference in sensitivity, the IC50 for venetoclax against both of the OSU-CLL lines was lower than the reported steady state plasma concentration for the drug (1.14 µM) when administered to CLL patients with deletion 17p at the standard 400mg dose (Salem, *et al* 2017).

Taken together, these observations suggest TR-57 may be effective at targeting CLL cells in the tumour microenvironment by interrupting key pro-survival signalling pathways and may represent an effective treatment option for high-risk CLL disease.

The increased expression of ATF4, Grp78 and HSP60 (Figure 4A) in CLL cells co-cultured with stromal cells, highlights these proteins as potentially important in promoting CLL cell survival in the tumour microenvironment. While the marked increase in ATF4 expression we observed following treatment with TR-57 may appear contradictory, our data are consistent with those of Ishizawa *et al*., who demonstrated that ONC-201 induces atypical activation of the UPR and cell death associated with increased expression of ATF4 in mantle cell lymphoma (MCL) and acute myeloid leukaemia (AML) cells (Ishizawa, *et al* 2016). Other studies also suggest ATF4 may function to promote both cell survival and apoptosis (Blais, *et al* 2004, Wortel, *et al* 2017) and that the differential roles of ATF4 are regulated by the formation of different ATF4 heterodimers (Kilberg, *et al* 2009). We also observed consistent down-regulation of heat shock protein 60 (HSP60) expression following TR-57 treatment (Figure 4A). Heat shock proteins play key roles in the adaption of cancer cells to ER stress and have been proposed as therapeutic targets for a range of cancers (Hsu, *et al* 2011, Lawson, *et al* 2015, Yun, *et al* 2020) including CLL (Frezzato, *et al* 2019, Guo, *et al* 2017). HSP60 has important roles in regulating cellular stress (Hall and Martinus 2013), mitochondrial dysfunction (Wu, *et al* 2017) and tumorigenesis (Merendino, *et al* 2010) and has been shown to act as both a suppressor as well as a tumour promoter in multiple cancers (Yun, *et al* 2020). A correlation between increased HSP60 expression and metastasis has also been observed in ovarian cancer (Guo, *et al* 2019), glioblastoma (Tang, *et al* 2016) and leukaemia (Wiechmann, *et al* 2017). The upregulation of HSP60 we observed in CLL cells co-cultured with CD40L-fibroblasts and the reduction in expression following treatment with TR-57 (Figure 4A) supports the notion that regulation of protein trafficking at the ER plays a crucial role in CLL-cell survival. Furthermore, the effects of TR-57 on HSP60 expression illustrates a previously undescribed mechanism by which drugs in this class may target protein homeostasis and induce apoptosis in cancer cells. The UPR plays an important role in the adaption of cancer cells to the stress associated with rapid cellular proliferation under nutrient and oxygen-deprived conditions. Emerging evidence suggests that a UPR also occurs within the mitochondrial proteome (UPR^mit^) that regulates cellular energy metabolism in both healthy and malignant cells (Deng and Haynes 2017). It is conceivable that TR-57 may target the UPR^mit^ in cancer cells by regulating activity of CIpP (Ishizawa, *et al* 2019), which leads to proteolysis of respiratory chain complexes and elevated levels of reactive oxygen species (ROS) (Zhang, *et al* 2020). In the current study, we showed that increased expression of CIpP following treatment with TR-57, alone and in combination with venetoclax (Figure 4A), was associated with down-regulation of MCL-1 and BCL-xL (Figure 4C). While it is unclear whether CIpP has any direct proteolytic activity towards BCL-2 family proteins, previous studies suggest increased ATF4 expression may lead to transcriptional upregulation of members of the BCL-2 family (Hetz 2012). This may explain the increased levels of NOXA and BAX we observed in CLL cells treated with TR-57 alone, which resulted in significant increases in the NOXA/MCL-1 and BAX/BCL-2 ratios (Figure 4C). However, it is unclear why treatment with TR-57 and venetoclax in combination lead to markedly lower levels of NOXA and BAX. The obvious explanation that this was due to excessive cell death following treatment with both drugs was excluded by assessing cell viability prior to cell lysis.

The increase in MCL-1 expression we observed following treatment with venetoclax (Figure 4C), may be significant in the sensitivity of CLL cells to this drug. This is supported by a recent study, which demonstrated MCL-1 plays a significant role in the resistance of AML cells to venetoclax (Carter, *et al* 2020). The data presented here suggest that synergy between TR-57 and venetoclax may be due to downregulation of MCL-1 and BCL-xL (Figure 4C), which raises the possibility that TR-57 may have therapeutic potential for CLL and AML patients who develop venetoclax-resistant disease. The effects of the pan caspase inhibitor, z-VAD-FMK, on cell death indicate that TR-57 induces caspase-dependent apoptosis (Supplementary Figure 2A). Interestingly, however z-VAD-FMK had no effect on cell death induced by venetoclax, alone or in combination with TR-57. Although further studies are required to confirm this under the conditions employed in the current study, evidence in the literature suggests that venetoclax may induce autophagy-mediated cell death in CLL cells by targeting the BCL-2/Beclin-2 complex (Avsec, *et al* 2021). If this is the case, synergy between TR-57 and venetoclax may be due to induction of both apoptosis and autophagy-mediated cell death pathways.

The effects of TR-57 on stroma-induced phosphorylation of AKT and ERK1/2-MAPK (Figure 4B) may also play a significant role in the cytotoxic effects of this drug against CLL cells. AKT plays a crucial role in CLL-cell survival and inhibition of ERK1/2-MAPK signalling downstream of the B-cell receptor is a key determinant of ibrutinib efficacy (Cheng, *et al* 2014). It is conceivable, but remains to be tested, whether TR-57 is effective against CLL cells with mutations of *BTK* or *PLC-γ2*, that confer resistance to ibrutinib (Woyach, *et al* 2014).

Collectively, the effects of TR-57 on signalling downstream of the BCR, protein trafficking and homeostasis and on the balance of expression of the BCL-2 family proteins demonstrate a multi-faceted mechanism of action of this drug against CLL cells. The cytotoxic and cytostatic synergy of TR-57 with concentrations of venetoclax below the reported steady state plasma concentrations (Salem, *et al* 2017), suggest that the benefits of the drug combination may be achieved *in vivo* with reduced doses of the BH3-mimetic. Furthermore, the effects of the drugs in combination on the proliferative and migratory capacities of CLL cells also raises the possibility that this drug combination may be effective against CLL cells in the tumour microenvironment and may represent a treatment option for patients with poor risk features, including those with TP53 aberrations.

## Acknowledgments

O.G.B., S.P.M., R.I.C., E.J.I., H.L. and D.S.K. designed the study and wrote the manuscript. N.F., Y.S., K.C. and O.G.B. generated and analysed the data.

## Figure and Table Legends

**Supplementary Figure 1.**
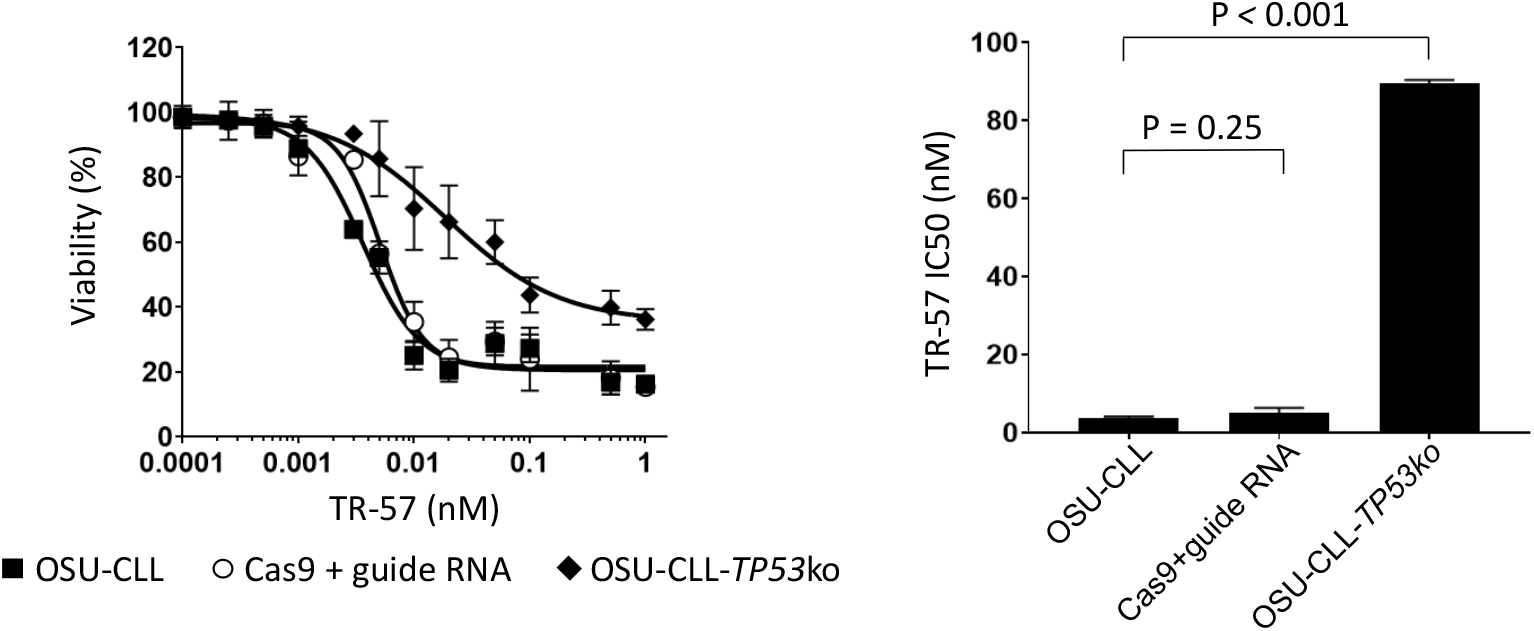
The transfection processes involved in generating the OSU-CLL-*TP53*ko cell line had no effect on the sensitivity of the OSU-CLL cells to TR-57. Dose response analysis of TR-57 against the OSU-CLL, OSU-CLL transfected with Cas9 and guide RNA and OSU-CLL-*TP53*ko lines. The cell lines were treated with the indicated doses of TR-57 for 48 h. Viability was assessed by flow cytometry and DiIC_1_(5) and propidium iodide staining. IC50 values were calculated from the dose-response analyses using GraphPad Prism software.

**Supplementary Table 1.**
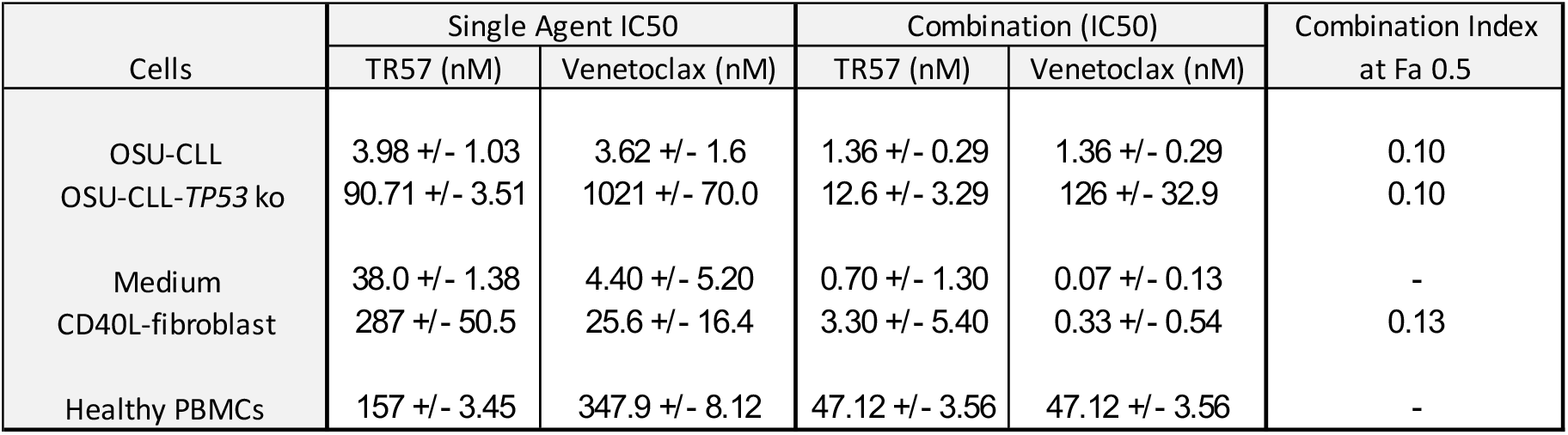
IC50 values for TR-57 and venetoclax, as single agents or in combination against CLL cells and PBMCs from healthy donors. IC50 values for TR-57 and venetoclax were calculated from dose-response analyses performed using flow cytometry and DiIC_1_(5) and propidium iodide following treatment for 48 h. Values shown are +/- standard deviation of at least 4 replicates or patient samples. Combination indices at a fractional effect (Fe) of 0.5 (50% cell killing) were calculated using CompuSyn software.

**Supplementary Table 2.** Combination indices (CI) for TR-57 and venetoclax in combination over a range of different fractional effect levels. The dose ranges, resulting fractional effects (Fe) and the combination indices (CI) for TR-57 and venetoclax in the OSU-CLL, OSU-CLL-*TP53*ko and CLL patient samples. The ratio of the drugs in combination were determined by the IC50s of the drugs as single agents in each cell type. Fractional effects (Fe) indicate the degree of cell killing, such that a Fe of 0.5 indicates 50% cell killing. CI values were calculated using CompuSyn software; values of <1 are indicative of synergy.

| OSU-CLL |  |  |  | OSU-CLL-TP53 ko |  |  |  | Primary CLL Cells |  |  |  |
| --- | --- | --- | --- | --- | --- | --- | --- | --- | --- | --- | --- |
| TR-57 (nM) | Venetoclax (nM) | Fe | Cl | TR-57 (nM) | Venetoclax (nM) | Fe | Cl | TR-57 (nM) | Venetoclax (nM) | Fe | Cl |
| 0.25 | 0.25 | 0.30 | 0.68 | 0.25 | 2.5 | 0.06 | 41.04 | 2.5 | 0.25 | 0.19 | 0.19 |
| 0.5 | 0.5 | 0.40 | 0.37 | 0.5 | 5 | 0.11 | 9.04 | 5 | 0.5 | 0.30 | 0.15 |
| 1.5 | 1.5 | 0.62 | 0.09 | 1.5 | 15 | 0.28 | 0.32 | 15 | 1.5 | 0.48 | 0.13 |
| 2.5 | 2.5 | 0.64 | 0.12 | 2.5 | 25 | 0.34 | 0.21 | 25 | 2.5 | 0.59 | 0.10 |
| 5 | 5 | 0.83 | 0.02 | 5 | 50 | 0.48 | 0.06 | 50 | 5 | 0.64 | 0.14 |
| 25 | 25 | 0.94 | 0.00 | 25 | 250 | 0.77 | 0.01 | 250 | 25 | 0.73 | 0.35 |
| 50 | 50 | 0.95 | 0.00 | 50 | 500 | 0.80 | 0.02 | 500 | 50 | 0.94 | 0.04 |
| 250 | 250 | 0.95 | 0.02 | 250 | 2500 | 0.82 | 0.07 | 2500 | 250 | 0.95 | 0.15 |
| 500 | 500 | 0.96 | 0.02 | 500 | 5000 | 0.89 | 0.04 | 5000 | 500 | 0.96 | 0.19 |

**Supplementary Figure 2.**
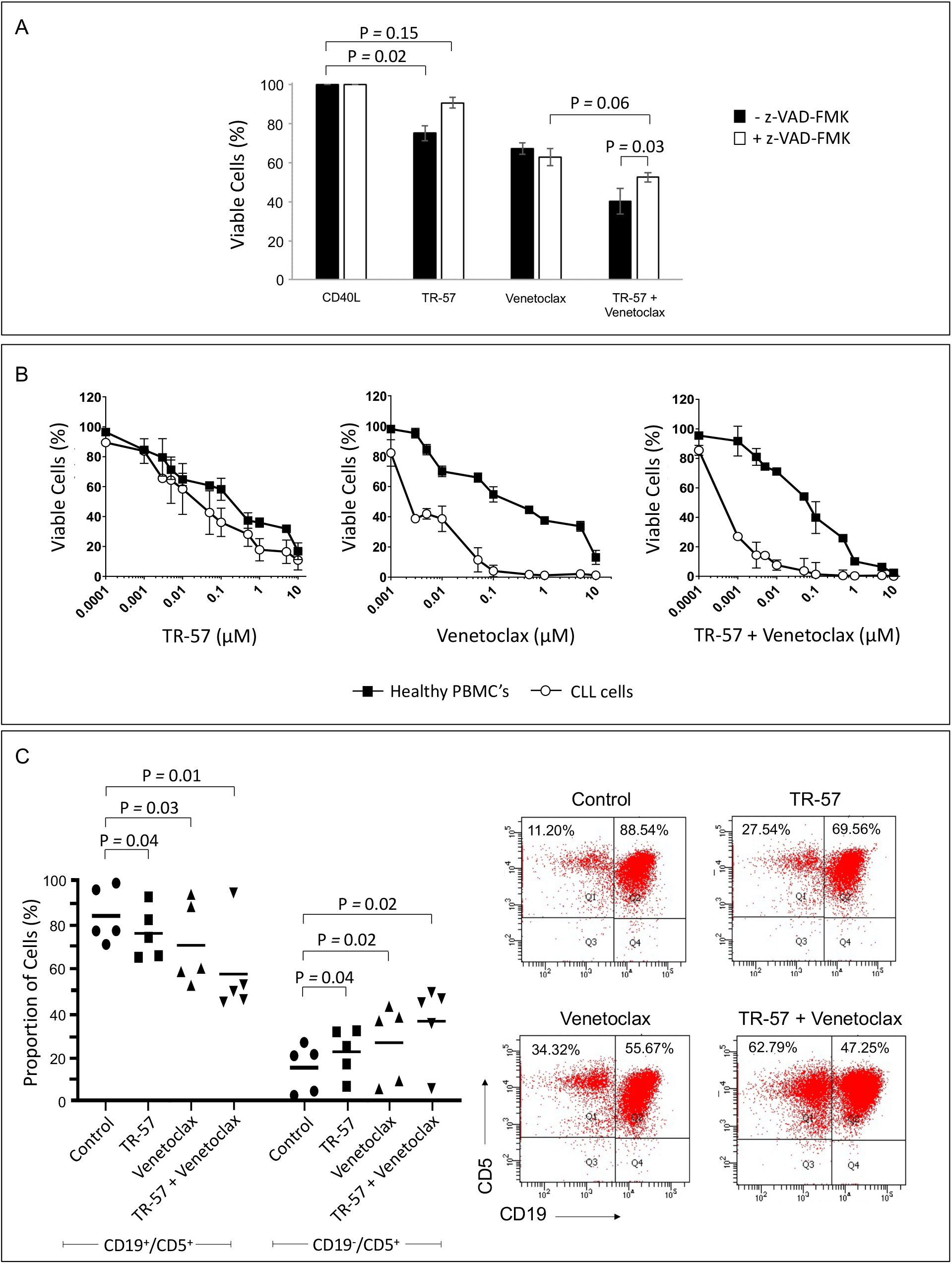
TR-57 induces caspase-dependent cell death of CLL cells and is more effective towards CLL cells than healthy PBMCs. (A) CLL patient samples (n = 3) were cultured with CD40L-fibroblasts in the presence or absence of the caspase inhibitor z-VAD-FMK (100 µM) for 1 h before being treated with 100 nM TR-57, 100 nM venetoclax, alone or in combination, for 48 h. Cell viability was assessed using DiIC_1_(5), propidium iodide and flow cytometry. (B) Dose-response analyses comparing the response of PBMCs and CLL cells to TR-57 and venetoclax, alone and in combination. Cell viability was assessed by flow cytometry using DiIC_1_(5) and propidium iodide. (B) Flow cytometry, DiIC_1_(5), propidium iodide and antibodies to CD19 and CD5 were used to assess the proportion of viable CLL and T-cells in PBMC fractions from CLL patients following treatment with TR-57 (100 nM) and venetoclax (10 nM), alone and in combination for 24 h.

**Supplementary Figure 3.**
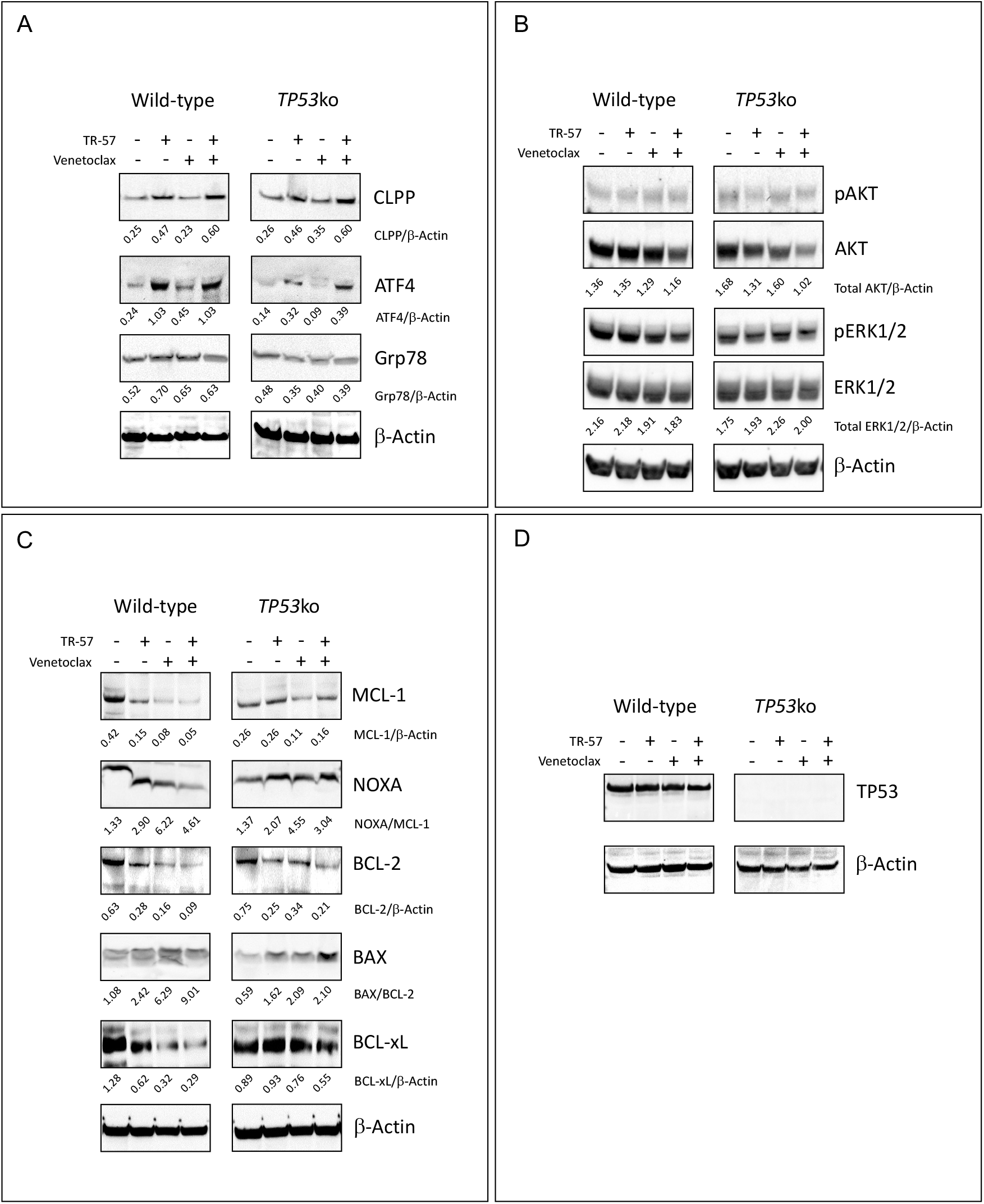
TR-57, alone and in combination with venetoclax, induces expression of CIpP and ATF-4 and a pro-apoptotic shift in the expression of BCL-2 family proteins in the OSU-CLL and OSU-CLL*TP53*ko cell lines. Immunoblotting for (A) UPR, (B) AKT and ERK1/2-MAPK, (C) BCL-2 family and (D) TP53 proteins in lysates from the OSU-CLL and OSU-CLL*TP53*ko cell lines following treatment with 5 nM TR-57 and 5 nM venetoclax (OSU-CLL) and 50 nM TR-57 and 500 nM venetoclax (OSU-CLL*TP53*ko), either alone or in combination for 24 h.

## Notes

### Competing Interest Statement

The authors have declared no competing interest.

